## Supplemental Information 3 for "An unclean slate, discrepancies between food input and recovered protein signal from experimental foodcrusts": Task1CalibrationTaskconfig.toml

```
TaskType = "Calibrate"

[CalibrationParameters]
WriteIntermediateFiles = false
WriteIndexedMzml = true
MinMS1IsotopicPeaksNeededForConfirmedIdentification = 3
MinMS2IsotopicPeaksNeededForConfirmedIdentification = 2
NumFragmentsNeededForEveryIdentification = 10

[CommonParameters]
TaskDescriptor = "Calibration_JD1"
MaxThreadsToUsePerFile = 20
ListOfModsFixed = "Common Fixed\tCarbamidomethyl on C\t\tCommon Fixed\tCarbamidomethyl on U"
ListOfModsVariable = "Common Variable\tOxidation on M"
DoPrecursorDeconvolution = true
UseProvidedPrecursorInfo = true
DeconvolutionIntensityRatio = 3.0
DeconvolutionMaxAssumedChargeState = 12
DeconvolutionMassTolerance = "±4.0000 PPM"
TotalPartitions = 1
ProductMassTolerance = "±25.0000 PPM"
PrecursorMassTolerance = "±15.0000 PPM"
AddCompIons = false
ScoreCutoff = 5.0
ReportAllAmbiguity = true
NumberOfPeaksToKeepPerWindow = 200
MinimumAllowedIntensityRatioToBasePeak = 0.01
NormalizePeaksAccrossAllWindows = false
TrimMs1Peaks = false
TrimMsMsPeaks = true
UseDeltaScore = false
QValueOutputFilter = 1.0
CustomIons = []
AssumeOrphanPeaksAreZ1Fragments = true
MaxHeterozygousVariants = 4
MinVariantDepth = 1
DissociationType = "HCD"
SeparationType = "HPLC"
MS2ChildScanDissociationType = "Unknown"
MS3ChildScanDissociationType = "Unknown"

[CommonParameters.DigestionParams]
MaxMissedCleavages = 2
InitiatorMethionineBehavior = "Variable"
MinPeptideLength = 7
MaxPeptideLength = 2147483647
MaxModificationIsoforms = 1024
MaxModsForPeptide = 2
Protease = "trypsin"
SearchModeType = "Full"
FragmentationTerminus = "Both"
SpecificProtease = "trypsin"
GeneratehUnlabeledProteinsForSilac = true
KeepNGlycopeptide = false
KeepOGlycopeptide = false

[[FileSpecificParameters]]
```
