## Supplemental Information 3 for "An unclean slate, discrepancies between food input and recovered protein signal from experimental foodcrusts": Task2GptmdTaskconfig.toml

```
TaskType = "Gptmd"

[GptmdParameters]
ListOfModsGptmd = "Common Biological\tAcetylation on K\t\tCommon Biological\tAcetylation on X\t\tCommon Biological\tADP-ribosylation on S\t\tCommon Biological\tButyrylation on K\t\tCommon Biological\tCarboxylation on D\t\tCommon Biological\tCarboxylation on E\t\tCommon Biological\tCarboxylation on K\t\tCommon Biological\tCitrullination on R\t\tCommon Biological\tCrotonylation on K\t\tCommon Biological\tDimethylation on K\t\tCommon Biological\tDimethylation on R\t\tCommon Biological\tFormylation on K\t\tCommon Biological\tGlu to PyroGlu on Q\t\tCommon Biological\tGlutarylation on K\t\tCommon Biological\tHexNAc on Nxs\t\tCommon Biological\tHexNAc on Nxt\t\tCommon Biological\tHexNAc on S\t\tCommon Biological\tHexNAc on T\t\tCommon Biological\tHydroxybutyrylation on K\t\tCommon Biological\tHydroxylation on K\t\tCommon Biological\tHydroxylation on N\t\tCommon Biological\tHydroxylation on P\t\tCommon Biological\tMalonylation on K\t\tCommon Biological\tMethylation on K\t\tCommon Biological\tMethylation on R\t\tCommon Biological\tNitrosylation on C\t\tCommon Biological\tNitrosylation on Y\t\tCommon Biological\tPhosphorylation on S\t\tCommon Biological\tPhosphorylation on T\t\tCommon Biological\tPhosphorylation on Y\t\tCommon Biological\tPyridoxal phosphate on K\t\tCommon Biological\tSuccinylation on K\t\tCommon Biological\tSulfonation on Y\t\tCommon Biological\tTrimethylation on K\t\tCommon Artifact\tAmmonia loss on C\t\tCommon Artifact\tAmmonia loss on N\t\tCommon Artifact\tCarbamyl on C\t\tCommon Artifact\tCarbamyl on K\t\tCommon Artifact\tCarbamyl on M\t\tCommon Artifact\tCarbamyl on R\t\tCommon Artifact\tCarbamyl on X\t\tCommon Artifact\tDeamidation on N\t\tCommon Artifact\tDeamidation on Q\t\tCommon Artifact\tWater Loss on E\t\tMetal\tCalcium on D\t\tMetal\tCalcium on E\t\tMetal\tCu[I] on D\t\tMetal\tCu[I] on E\t\tMetal\tFe[II] on D\t\tMetal\tFe[II] on E\t\tMetal\tFe[III] on D\t\tMetal\tFe[III] on E\t\tMetal\tMagnesium on D\t\tMetal\tMagnesium on E\t\tMetal\tPotassium on D\t\tMetal\tPotassium on E\t\tMetal\tSodium on D\t\tMetal\tSodium on E\t\tMetal\tZinc on D\t\tMetal\tZinc on E\t\tTrypsin Digested\tEQIGG (sumoylation (SMT-3) Site yeast) on K\t\tTrypsin Digested\tGG (Ubiquitination Site) on K"

[CommonParameters]
TaskDescriptor = "GPTMD_JD_ExpFC_02"
MaxThreadsToUsePerFile = 20
ListOfModsFixed = "Common Fixed\tCarbamidomethyl on C\t\tCommon Fixed\tCarbamidomethyl on U"
ListOfModsVariable = "Common Variable\tOxidation on M"
DoPrecursorDeconvolution = true
UseProvidedPrecursorInfo = true
DeconvolutionIntensityRatio = 3.0
DeconvolutionMaxAssumedChargeState = 12
DeconvolutionMassTolerance = "±4.0000 PPM"
TotalPartitions = 1
ProductMassTolerance = "±0.5000 Absolute"
PrecursorMassTolerance = "±3.0000 PPM"
AddCompIons = false
ScoreCutoff = 5.0
ReportAllAmbiguity = true
NumberOfPeaksToKeepPerWindow = 200
MinimumAllowedIntensityRatioToBasePeak = 0.01
NormalizePeaksAccrossAllWindows = false
TrimMs1Peaks = false
TrimMsMsPeaks = true
UseDeltaScore = false
QValueOutputFilter = 1.0
CustomIons = []
AssumeOrphanPeaksAreZ1Fragments = true
MaxHeterozygousVariants = 4
MinVariantDepth = 1
DissociationType = "HCD"
SeparationType = "HPLC"
MS2ChildScanDissociationType = "Unknown"
MS3ChildScanDissociationType = "Unknown"

[CommonParameters.DigestionParams]
MaxMissedCleavages = 2
InitiatorMethionineBehavior = "Variable"
MinPeptideLength = 7
MaxPeptideLength = 2147483647
MaxModificationIsoforms = 1024
MaxModsForPeptide = 2
Protease = "trypsin"
SearchModeType = "Full"
FragmentationTerminus = "Both"
SpecificProtease = "trypsin"
GeneratehUnlabeledProteinsForSilac = true
KeepNGlycopeptide = false
KeepOGlycopeptide = false
```
