## Supplemental Information 3 for "An unclean slate, discrepancies between food input and recovered protein signal from experimental foodcrusts": Task3SearchTaskconfig.toml

```
TaskType = "Search"

[SearchParameters]
DisposeOfFileWhenDone = true
DoParsimony = true
ModPeptidesAreDifferent = false
NoOneHitWonders = true
MatchBetweenRuns = false
Normalize = false
QuantifyPpmTol = 5.0
DoHistogramAnalysis = false
SearchTarget = true
DecoyType = "Reverse"
MassDiffAcceptorType = "OneMM"
WritePrunedDatabase = false
KeepAllUniprotMods = true
DoLocalizationAnalysis = true
DoQuantification = true
SearchType = "Classic"
LocalFdrCategories = ["FullySpecific"]
MaxFragmentSize = 30000.0
MinAllowedInternalFragmentLength = 0
HistogramBinTolInDaltons = 0.003
MaximumMassThatFragmentIonScoreIsDoubled = 0.0
WriteMzId = true
WritePepXml = false
WriteDecoys = true
WriteContaminants = true
WriteIndividualFiles = true
WriteSpectralLibrary = false
CompressIndividualFiles = false
TCAmbiguity = "RemoveContaminant"
IncludeModMotifInMzid = false

[SearchParameters.ModsToWriteSelection]
'N-linked glycosylation' = 3
'O-linked glycosylation' = 3
'Other glycosylation' = 3
'Common Biological' = 3
'Less Common' = 3
Metal = 3
'2+ nucleotide substitution' = 3
'1 nucleotide substitution' = 3
UniProt = 2

[CommonParameters]
TaskDescriptor = "Search_JD_ExpFC_02"
MaxThreadsToUsePerFile = 20
ListOfModsFixed = "Common Fixed\tCarbamidomethyl on C\t\tCommon Fixed\tCarbamidomethyl on U"
ListOfModsVariable = "Common Variable\tOxidation on M"
DoPrecursorDeconvolution = true
UseProvidedPrecursorInfo = true
DeconvolutionIntensityRatio = 3.0
DeconvolutionMaxAssumedChargeState = 12
DeconvolutionMassTolerance = "±4.0000 PPM"
TotalPartitions = 1
ProductMassTolerance = "±0.5000 Absolute"
PrecursorMassTolerance = "±3.0000 PPM"
AddCompIons = false
ScoreCutoff = 5.0
ReportAllAmbiguity = true
NumberOfPeaksToKeepPerWindow = 200
MinimumAllowedIntensityRatioToBasePeak = 0.01
NormalizePeaksAccrossAllWindows = false
TrimMs1Peaks = false
TrimMsMsPeaks = true
UseDeltaScore = false
QValueOutputFilter = 1.0
CustomIons = []
AssumeOrphanPeaksAreZ1Fragments = true
MaxHeterozygousVariants = 4
MinVariantDepth = 1
DissociationType = "HCD"
SeparationType = "HPLC"
MS2ChildScanDissociationType = "Unknown"
MS3ChildScanDissociationType = "Unknown"

[CommonParameters.DigestionParams]
MaxMissedCleavages = 2
InitiatorMethionineBehavior = "Variable"
MinPeptideLength = 7
MaxPeptideLength = 2147483647
MaxModificationIsoforms = 1024
MaxModsForPeptide = 2
Protease = "trypsin"
SearchModeType = "Full"
FragmentationTerminus = "Both"
SpecificProtease = "trypsin"
GeneratehUnlabeledProteinsForSilac = true
KeepNGlycopeptide = false
KeepOGlycopeptide = false
```
